# Learning Drug Function from Chemical Structure with Convolutional Neural Networks and Random Forests

**DOI:** 10.1101/482877

**Authors:** Jesse G. Meyer, Shengchao Liu, Ian J. Miller, Joshua J. Coon, Anthony Gitter

## Abstract

Empirical testing of chemicals for drug efficacy costs many billions of dollars every year. The ability to predict the action of molecules *in silico* would greatly increase the speed and decrease the cost of prioritizing drug leads. Here, we asked whether drug function, defined as MeSH “Therapeutic Use” classes, can be predicted from only chemical structure. We evaluated two chemical structure-derived drug classification methods, chemical images with convolutional neural networks and molecular fingerprints with random forests, both of which outperformed previous predictions that used drug-induced transcriptomic changes as chemical representations. This suggests that a chemical’s structure contains at least as much information about its therapeutic use as the transcriptional cellular response to that chemical. Further, because training data based on chemical structure is not limited to a small set of molecules for which transcriptomic measurements are available, our strategy can leverage more training data to significantly improve predictive accuracy to 83-88%. Finally, we explore use of these models for prediction of side effects and drug repurposing opportunities, and demonstrate the effectiveness of this modeling strategy for multi-label classification.

## Introduction

Development of molecules with new or improved properties is needed in many industries, including energy, agriculture, and medicine. However, the number of possible molecules to explore, also referred to as chemical space, is exceedingly large^1,2^. Even when chemical space is limited to compounds that conform to ‘Lipinski’s Rule of Five’^3^, which applies to the subtask of drug development, there are still as many as 10^60^ possible chemical structures^4^. Regardless of the available chemical diversity, the pace of new drug approvals has steadily decreased, leaving room for new approaches that can improve the current process.

A promising approach for discovering new drug molecules is machine learning^5,6^, which includes so-called deep learning using deep neural networks (DNNs)^7^. Many studies describe methods for embedding molecules into a latent space and engineering molecules with desirable properties^8–10^. Reinforcement learning has been applied with paired DNNs to design molecules with desired properties, such as solubility or transcription factor inhibition^11^. A framework for benchmarking model predictions is available^12^. One study used a generative adversarial neural network architecture to generate molecules that should induce specific transcriptomic states^13^. There are many ways to represent molecules for machine learning. Many papers use SMILES strings^14–16^ as molecular inputs for embedding, but there is a trend toward use of molecular graphs^17–19^.

A general weakness of DNNs is that they perform best with large amounts of training data (100,000 to millions of examples, e.g. ImageNet^20^). However, DNNs can be used for problems with small training data through transfer learning, where networks are trained on a large dataset for one problem and adapted for a related problem that has less training data ^21–23^. For example, transfer learning has been applied to classification of less than 6,000 medical ultrasound images^24^, only 2,000 oceanfront images^25^, or less than 1,000 cellular images^26^, even though these networks were pretrained on images of completely different objects.

One type of DNN for structured data such as sequences and images, the convolutional neural network (CNN), has enabled major advances in image processing tasks in diverse fields. In chemistry, several papers have described excellent performance resulting from use of two-dimensional images of chemicals with CNNs. This approach has been used effectively to predict chemical toxicity^27^ with regard to the 12 biological toxicity endpoints in the Tox21 challenge^28^. CNNs for chemical images have also been described as a general-purpose molecule property prediction tool despite their lack explicit chemistry knowledge^29^. These authors found that augmenting the same deep learning architecture with only three additional chemical properties further improved model performance^30^, suggesting that chemical images alone may not entirely capture the important characteristics of a chemical. Finally, images and CNNs have been used to predict drug-protein interactions and outperformed models trained on flattened versions of the images, which cannot exploit the spatial structure^31^.

The various ways to measure and represent molecules leads to a philosophical question about the nature of chemicals^32^, and the related fundamental question of whether a single representation can completely describe a chemical entity (reviewed in ^9^). As described above, several chemical structure-derived embeddings are often used for cheminformatics, such as chemical images or circular molecular fingerprints. Alternatively, molecules can be represented by an analytical measurement^33^ or by their influence on biological systems (e.g. transcriptomic or morphological changes)^34,35^. There are open questions regarding the relative utility of these various chemical representation strategies for different predictive tasks relevant to drug development.

In this paper, we use machine learning and chemical structure-derived molecule representations to predict specific medical subheading (MeSH) ‘Therapeutic Uses’ classes^36^. We first performed the same classification task with the same set of 676 molecules previously selected by Aliper *et al.*^34^. In contrast to our chemical structure-derived models, Aliper *et al.* used molecule-induced transcriptome changes from the LINCS project^37^ as a proxy molecule representation. We employed two strategies: (1) chemical images with CNNs, and (2) Morgan fingerprints (MFP)^38^ with random forests (RF)^39^ (**Figure 1**). We chose to use the CNN with images because of extensive precedent for the effectiveness of this pair, and we chose to use MFP with RF because we and others have seen excellent performance of this representation-model pair^40^. Our goal was to assess whether drug function classifier models trained with readily-available chemical structure input can outperform models trained with empirical measures of drug effects. Our results support the effectiveness of chemical structure-based models. Both classification models trained with chemical structure-derived features greatly outperform the previous benchmark based on drug-induced transcriptomic changes. Further, because we only require chemical structure, the models can be greatly improved by training on over 6,000 additional compounds that do not have associated transcriptomic data. Our main contribution is that chemical structures alone are effective predictors of therapeutic use classes.

**Figure 1:**
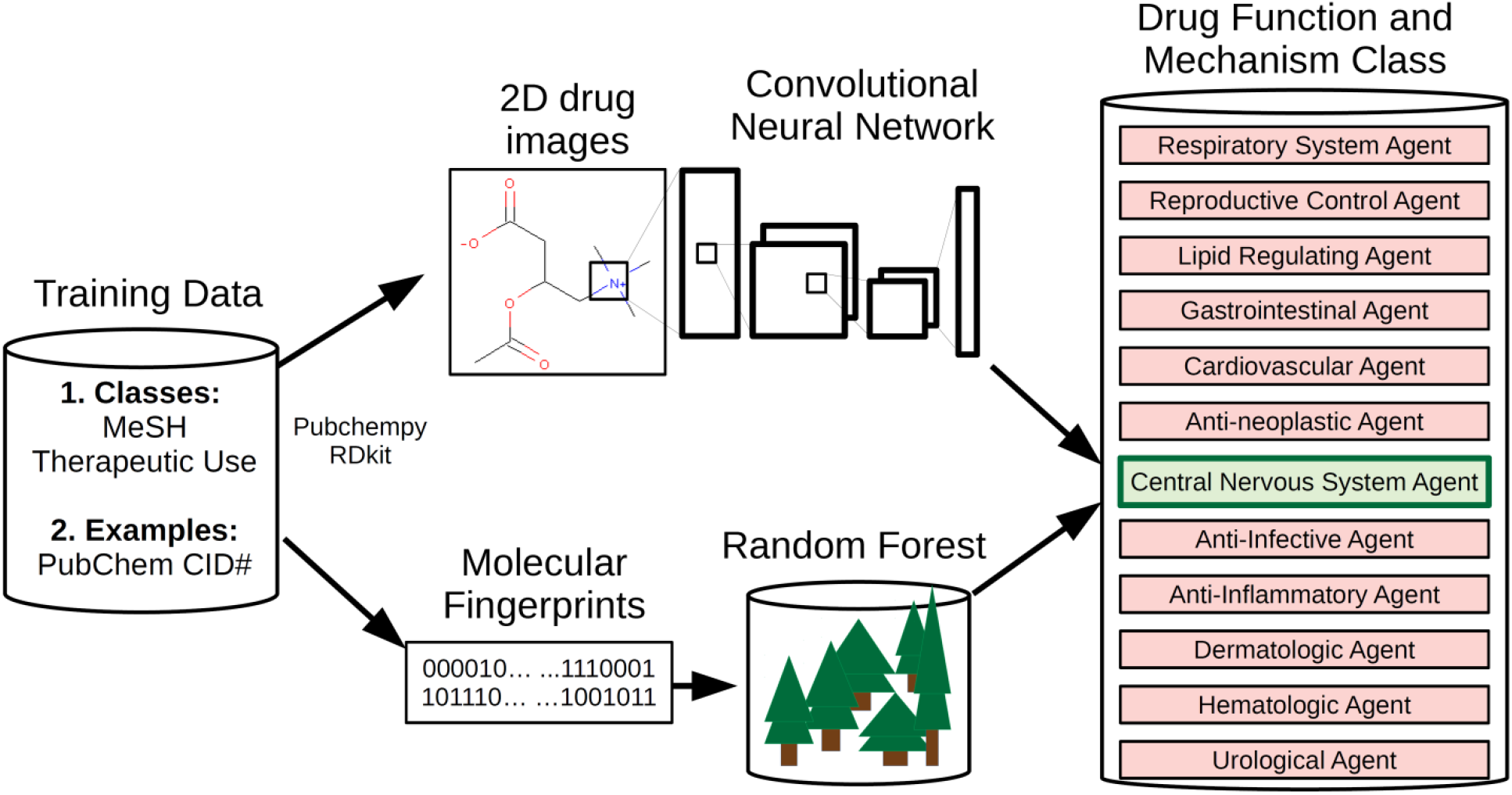
Structure-based drug classification pipelines. Chemicals from 12 medical subheadings (MeSH) therapeutic use classifications were converted to either 2-dimensional color molecule images or Morgan molecular fingerprints. Molecule images were used to train a convolutional neural network (IMG+CNN) classifier, and fingerprints were used to train a random forest (MFP+RF) classifier. The models were used separately to predict classes of drugs using stratified cross validation.

## METHODS

All code for the CNN and RF models and the pre-processed datasets are available from https://github.com/jgmeyerucsd/drug-class.

### Data

The primary goal of this work was to compare empirically-derived chemical features, such as the transcriptome-based model from Aliper *et al*.^34^, with chemical structure-derived representations. Therefore, in the first evaluation, we emulated their framing of the prediction task, which is to predict 1 of 12 MeSH ‘Therapeutic Use’ classes of chemicals. The specific version of the data used by Aliper *et al.*, including training/validation groups, is unavailable. Therefore, our exact training and validation sets are different. To make the fairest-possible comparison, we constructed a dataset following the same guidelines as Aliper *et al*.

Molecules were selected from PubChem^41^ on October 2nd, 2018 according to their MeSH ‘Therapeutic Uses’ classification (Chemicals and Drugs Category > Chemical Actions and Uses > Pharmacologic Actions > Therapeutic Uses). Although there are 20 high-level categories, we only used the 12 classes described previously^34^. Molecules in these 12 classes were downloaded in a spreadsheet containing their compound identification number (CID). A total of 11,929 CIDs were converted to SMILES strings using the Python package *pubchempy* (https://github.com/mcs07/PubChemPy). SMILES strings with length over 400, or membership to more than 1 of 12 MeSH therapeutic classes were excluded, leaving 8,372 SMILES. For this analysis, chemicals in multiple classes were excluded as described previously to enable direct comparison^34^. This final list was filtered to remove multiple versions of molecules that differ by only accompanying salts. The final filtered total was 6,955 molecules. The distribution of molecules among classes is given in **Table 1.** This set of all molecules was divided into five folds stratified based on class for cross validation.

SMILES strings were converted to three-color (RGB) images with size 500 x 500 pixels or 1024-bit Morgan fingerprints using the python package *RDKit*^42^. Images generated by RDKit always fit the entire molecule structure, so molecules of different sizes are not problematic. All images used for training and validation are available on GitHub. Molecule classes were then split into three subgroups for model training and prediction: 3-, 5-, or 12-class prediction tasks according to the groupings described previously by Aliper *et al.* (**Table 1**).

For the comparison with Aliper *et al.*’s results, we took the list of molecules in their supplemental table 1 and retrieved SMILES strings from PubChem using pubchempy. Images were generated as described above. During the removal of salts from their original set of 678 molecules we found that two drugs, one from anti-infective and one from CNS, were the same molecule with different salt pairs. The copy of these two duplicate molecules were removed. The numbers of chemicals in this smaller dataset are given in **Table 1**. This set of 676 molecules was split into 10 folds stratified based on class membership to mimic the methods or Aliper *et al.* as closely as possible. However, the dermatological and urological classes have less than 10 molecules and therefore are missing validation examples in some folds. For those folds missing validation examples, the receiver operator characteristic area under the curve (ROC AUC) and average precision metrics were not computed.

**Table 1:**
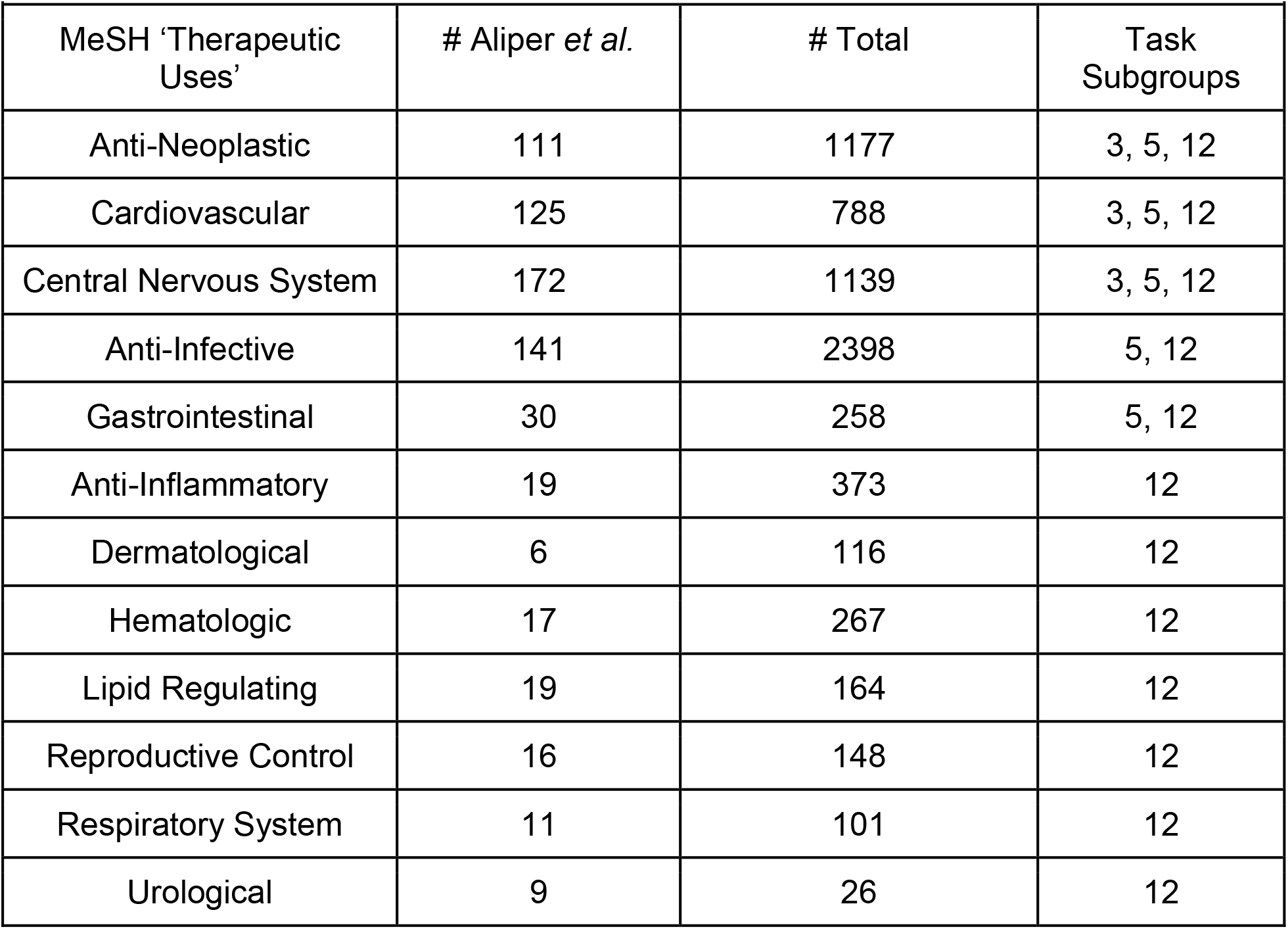
Summary of data classes and task groupings.

| **Table 1: Summary of data classes and task groupings.** |  |  |  |
| --- | --- | --- | --- |
| MeSH 'Therapeutic Uses' | # Aliper *et al.* | # Total | Task Subgroups |
| Anti-Neoplastic | 111 | 1177 | 3, 5, 12 |
| Cardiovascular | 125 | 788 | 3, 5, 12 |
| Central Nervous System | 172 | 1139 | 3, 5, 12 |
| Anti-Infective | 141 | 2398 | 5, 12 |
| Gastrointestinal | 30 | 258 | 5, 12 |
| Anti-Inflammatory | 19 | 373 | 12 |
| Dermatological | 6 | 116 | 12 |
| Hematologic | 17 | 267 | 12 |
| Lipid Regulating | 19 | 164 | 12 |
| Reproductive Control | 16 | 148 | 12 |
| Respiratory System | 11 | 101 | 12 |
| Urological | 9 | 26 | 12 |

### Images with Convolutional Neural Networks (IMG+CNN) – Single Label

Molecule images with RGB channels were resized to 150×150 pixels and used for retraining and validation of a CNN with predetermined weights from resnext101_64^43^ implemented using fastai and pytorch^44^. The loss function used was binary cross entropy and the output layer was logsoftmax. A cyclic cosine annealing learning rate was used during training^45^, which decreases from the initial setting toward 0 over a number of epochs. The number of epochs needed to decay the learning rate to the final value was length was doubled every cycle. An example of the learning rate versus batch is shown in **Figure S1** along with the corresponding training loss.

To determine the best hyperparameters for all CNN models and data subsets, we first performed hyperparameter optimization on the small set of 678 compounds. Hyperparameter optimization was done with nested 10-fold cross validation using class-stratified folds. Varied hyperparameters were: (1) dropout proportions of 20%, 40% or 60%, (2) retraining all weights or only the output layer weights, and (3) the initial learning rates for cosine annealing ([5e-5, 4e-4, 3e-3] or [1e-4, 1e-3, 1e-2] for early, middle, and output layers). Fixed training hyperparameters were the batch size of 25, seven cycles of cosine annealing learning rate with decay rate decreased by half each cycle (totaling 127 epochs), and data augmentation with random zooms of up to 10% and random horizontal or vertical image flips. Average accuracy values from each of the hyperparameter groups tested during the inner loops of nested cross validation are given in **Table S1**. Based on the results of this hyperparameter search, the hyperparameters that most often resulted in the best accuracy on the inner loop fold were used for training all other models, including CNNs trained on the larger set of 6,955 compounds. The tested learning rates had a minimal effect on the accuracy. The largest effect on accuracy resulted from retraining all weights instead of training only the output weights. These best hyperparameters from the grid were (1) 40% dropout, (2) retraining all weights, and (3) the higher learning rate set of [1e-4, 1e-3, 1e-2] for early, middle, and output neuron layer groups, respectively.

### Molecular Fingerprints with Random Forests (MFP+RF)

Random forests^39^ are ensembles of decision trees, where each tree is learned on a subsample of data points and features (in this case, bits in a molecular fingerprint). Benchmarking studies often include MFP+RF models because they are easy to train and have strong performance on a variety of computational chemistry tasks^12,40,46–50^. The random forest model was implemented with scikit-learn^51^. Separate hyperparameter grid searches (216 combinations, **Table 2**) were performed for the 676 and 6,955 compound analyses in a nested cross validation setting. For each outer loop, the best set of hyperparameters was selected based on the inner loop cross validation accuracy. These hyperparameters were then used to train on all the inner loop compounds and assess performance on the outer loop validation set.

**Table 2:** Random Forest Classifier Hyperparameter Sets

| **Table 2: Random Forest Classifier Hyperparameter Sets** |  |
| --- | --- |
| Parameter | Values |
| # estimators | 50, 250, 1000, 4000, 8000, 16000 |
| Max features | None, Sqrt, Log2 |
| Min sample leaf | 1, 10, 100, 1000 |
| Class weight | None, balanced subsample, balanced |

### Comparison of Drug-like Properties

CIDs were used to download Molecular weight, XLogP, HBondAcceptorCount, HBondDonorCount, and IsomericSMILES values using pubchempy. Compounds were then filtered to include non-redundant IsomericSMILES values and only drugs with a single class label (**Table S2**). XLogP values are computed^52^ rather than measured, and were not available for all queried compounds (**Table S3**). Violin plots were created using ggplot2 (https://ggplot2.tidyverse.org/). For each quantitative feature, a Welch’s ANOVA and Games-Howell post hoc test (R package userfriendlyscience, https://cran.r-project.org/web/packages/userfriendlyscience/index.html) were used to compare differences between chemical features between drug-class groups. This test was selected because it does not assume a normal distribution, even variance, or equal sample sizes between groups^53^. Adjusted p-values from the Games-Howell post hoc test are reported in **Table S4**. Drug-class level distribution and pairwise relations of chemical features were further visualized with Seaborn (https://seaborn.pydata.org/).

### Single-Label Classification Models: Training, Comparison and Evaluation

Training data for 676 molecules with transcriptomic measurements available was split into 10 folds for cross validation, and the training data for all 6,955 available annotated molecules in the 12 MeSH classes was split into 5 folds. When referring to model performance and metrics, all values are from the held-out folds referred to as validation folds, and metrics are the average performance of the validation folds unless otherwise specified. We checked the chemical similarity of our 5 folds of the larger dataset using ChemTreeMap^54^ and found the folds to be randomly distributed in chemical space (**Figure S2**). Trained models were evaluated using the following metrics from scikit-learn 0.20.3: accuracy, balanced accuracy, Matthew’s correlation coefficient (MCC), ROC AUC, and average precision score. The classification accuracy on the five validation sets was used as the primary model comparison metric. Class-specific prediction accuracy of the models was compared using confusion matrices for a single representative validation fold. We also compared our prediction accuracy with the accuracy previously reported by Aliper *et al.*^34^. However, it should be clearly noted that although we used the same molecules, the exact training and validation sets were unavailable, so the accuracies are not perfectly comparable.

### IMG+CNN – Multi-label Classification

The set of all molecules including those with multiple class memberships (8,336 molecules assigned a total of 9,885 classes) was used to train additional convolutional neural networks for multi-label classification using the fastai package. The data was split into five folds based on pairwise class co-occurrence using the iterative class splitter from the skmultilearn package^55^. A Jupyter notebook containing the code used to train the models is available on the GitHub repository under multiclass_data/multiclass_5foldCV.ipynb. Resnet50 was used as the pretrained model and weights, and 40% dropout was used with image data augmentation. Training images were 256×256 and were processed in batches of 40. All weights were retrained for each CNN model for 127 epochs (the same number as for single class) using the updated one-cycle policy^56^.

The multi-label classification was evaluated by computing thresholded accuracy and F-beta (beta of 2.0, the fastai default) using a default score cutoff of 0.5. ROC AUC and average precision scores were also computed as described for the single-label classification models using the weighted average. Finally, a network of the class relationships was computed using the pairwise co-occurrence of classes using the networkx (https://networkx.github.io/) and igraph (https://igraph.org/python/) Python packages according to the skmultilearn tutorial (http://scikit.ml/labelrelations.html). Network graphs were visualized with their edge width proportional to the strength of the node relationship as defined by the number of co-occurrences of the classes.

## RESULTS

### Classification with Small Benchmark Dataset

Two chemical structure-derived representations were used for training and classification with two different model architectures: (1) IMG+CNN or (2) MFP+RF (**Figure 1**). Molecules were split into three subtask sets as described previously^34^. Each subtask set contained 408, 579, or 676 molecules for the 3-, 5- and 12-class problems, respectively. **Table 3** gives a summary of the validation set accuracy for the models described here in comparison with results from Aliper *et al*. who used a multilayer perceptron DNN or support vector machine (SVM) with gene expression changes as model input. For 5- and 12-class subtasks, MFP+RF performed best, achieving 64.1% accuracy on the 12-class prediction task, representing an improvement over the expression-based DNN that achieved only 54.6% accuracy. The IMG+CNN model produced accuracy similar to the MFP+RF for the 3-class subtask, but achieved about 4 percentage points worse accuracy on the 5- and 12-class problems. However, IMG+CNN models still significantly outperformed the previous gene expression-based models in all cases. Accuracy can be inflated when the class labels are not evenly distributed (**Table 1**). Balanced accuracy and MCC are more robust with skewed classes but were not reported for the gene expression-based models. For the IMG+CNN and MFP+RF, these values are lower than the accuracy but good overall.

**Table 3:** Average metrics for each of 10 hold-out folds from cross validation using 676 molecules from Aliper *et al.* annotated with only one of the 12 MeSH classes. Values for the gene expression-based models are from Aliper *et al.* who used different training and validation folds for 10-fold cross validation with a ^1^support vector machine (SVM) or ^2^multilayer perceptron deep neural network (DNN) based on pathway activation scores. Values from this paper using ^3^molecule images input to a convolutional neural network (IMG+CNN) or ^4^Morgan molecular fingerprints as input to the random forest (MFP+RF). Values for ^3,4^ are the mean of the validation folds ± standard deviation. *Area under the receiver operating characteristic (ROC AUC) and average precision score were computed as the weighted average of scores across classes and only computed for the first 6 validation sets of the 12-class problem due to less than 10 examples in the dermatological and urological classes.

| **Problem group** | **Metric** | **SVM<sup>1</sup>** | **DNN<sup>2</sup>** | **IMG+CNN<sup>3</sup>** | **MFP+RF<sup>4</sup>** |
| --- | --- | --- | --- | --- | --- |
| **3-class** | **Accuracy** | **0.53** | **0.701** | **$0.747 \pm 0.0657$** | **$0.742 \pm 0.0692$** |
|  | **Balanced Accuracy** |  |  | **$0.739 \pm 0.0644$** | **$0.715 \pm 0.0766$** |
|  | **MCC** |  |  | **$0.619 \pm 0.102$** | **$0.612 \pm 0.106$** |
|  | **ROC AUC** |  |  | **$0.870 \pm 0.0412$** | **$0.894 \pm 0.0417$** |
|  | **Ave. Precision Score** |  |  | **$0.806 \pm 0.0592$** | **$0.847 \pm 0.0588$** |
| **5-class** | **Accuracy** | **0.417** | **0.596** | **$0.653 \pm 0.0451$** | **$0.694 \pm 0.0497$** |
|  | **Balanced Accuracy** |  |  | **$0.620 \pm 0.0509$** | **$0.635 \pm 0.0661$** |
|  | **MCC** |  |  | **$0.549 \pm 0.0599$** | **$0.606 \pm 0.0660$** |
|  | **ROC AUC** |  |  | **$0.867 \pm 0.0322$** | **$0.892 \pm 0.0284$** |
|  | **Ave. Precision Score** |  |  | **$0.735 \pm 0.0568$** | **$0.791 \pm 0.0471$** |
| **12-class** | **Accuracy** | **0.366** | **0.546** | **$0.608 \pm 0.0500$** | **$0.641 \pm 0.0331$** |
|  | **Balanced Accuracy** |  |  | **$0.507 \pm 0.107$** | **$0.504 \pm 0.0522$** |
|  | **MCC** |  |  | **$0.525 \pm 0.0620$** | **$0.572 \pm 0.0388$** |
|  | **ROC AUC\*** |  |  | **$0.863 \pm 0.209$** | **$0.896 \pm 0.0200$** |
|  | **Ave. Precision Score\*** |  |  | **$0.672 \pm 0.0303$** | **$0.751 \pm 0.0205$** |

### Classification with All Annotated Chemicals

A major limitation of using empirical chemical features generated from biological experiments, such as gene expression, is that the time and cost limit the size of training data available for models. For the drug function prediction task, there are roughly 10x more molecules available annotated with the 12 MeSH classes than the number of molecules with transcriptomic data. To highlight the value of using a chemical-based representation instead of an empirical representation, we trained additional models with all available 6,955 chemical structures. The use of more training data was greatly beneficial to both representation-model pairs resulting in accuracies of 83-88% and ROC AUC values over 0.969 (**Table 4**). ROC curves for predictions from the IMG+CNN model are shown in **Figure 2**, and curves for the MFP+RF are shown in **Figure S3.** With this larger training dataset, the 5-fold cross validation evaluation metrics are quite similar for the IMG+CNN and MFP+RF models. The MFP+RF model has a slight advantage over the IMG+CNN model when using the metrics that consider the complete rankings of chemicals by predicted class probabilities (ROC AUC and average precision).

**Table 4:**
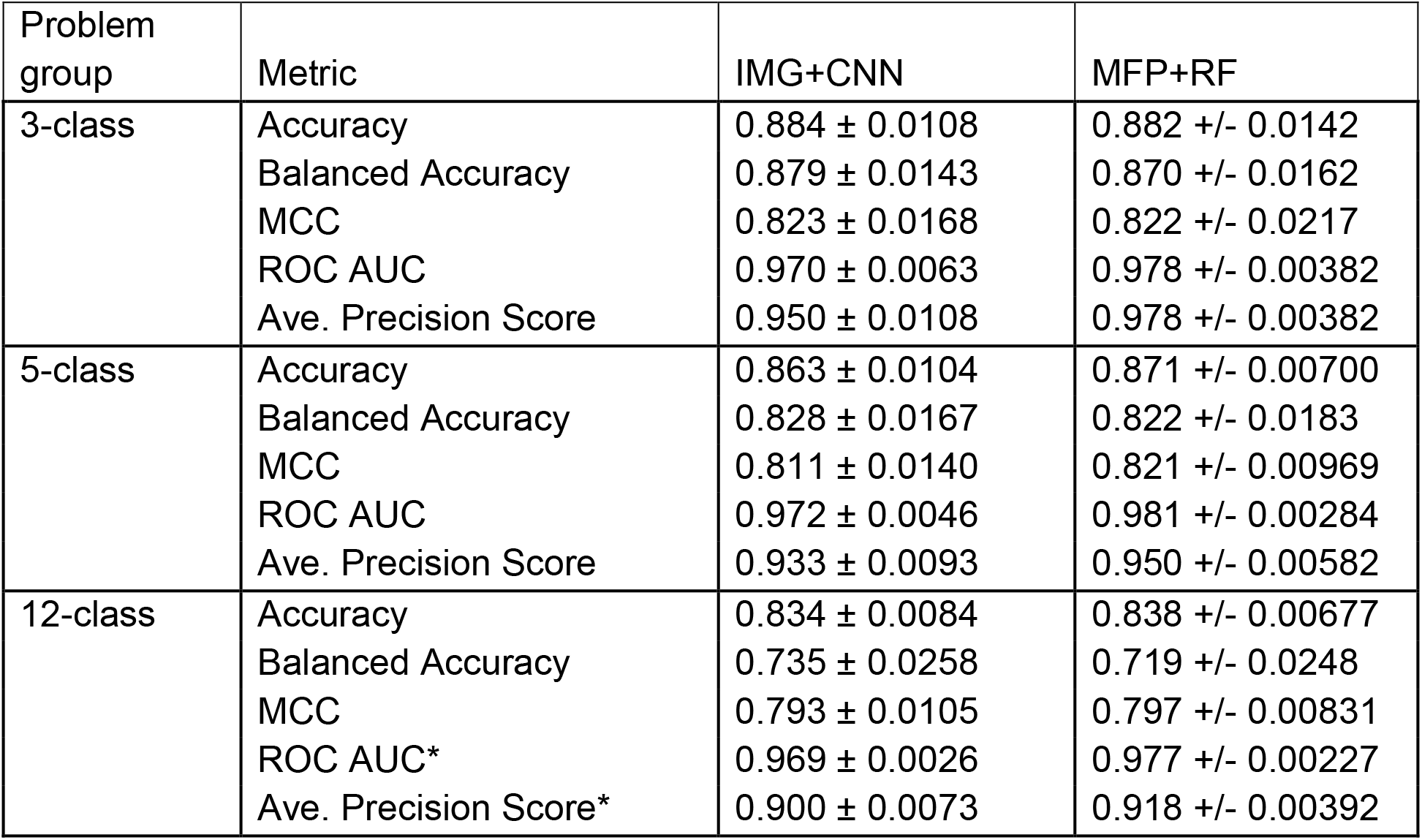
Average metrics for each of 5 validation folds from cross validation using the full set of 6,955 molecules annotated with only one of the 12 MeSH classes. *Area under the receiver operating characteristic (ROC AUC) and average precision score were computed as the weighted average of scores across classes.

| Problem group | Metric | IMG+CNN | MFP+RF |
| --- | --- | --- | --- |
| 3-class | Accuracy | 0.884 ± 0.0108 | 0.882 +/- 0.0142 |
| | Balanced Accuracy | 0.879 ± 0.0143 | 0.870 +/- 0.0162 |
| | MCC | 0.823 ± 0.0168 | 0.822 +/- 0.0217 |
| | ROC AUC | 0.970 ± 0.0063 | 0.978 +/- 0.00382 |
| | Ave. Precision Score | 0.950 ± 0.0108 | 0.978 +/- 0.00382 |
| 5-class | Accuracy | 0.863 ± 0.0104 | 0.871 +/- 0.00700 |
| | Balanced Accuracy | 0.828 ± 0.0167 | 0.822 +/- 0.0183 |
| | MCC | 0.811 ± 0.0140 | 0.821 +/- 0.00969 |
| | ROC AUC | 0.972 ± 0.0046 | 0.981 +/- 0.00284 |
| | Ave. Precision Score | 0.933 ± 0.0093 | 0.950 +/- 0.00582 |
| 12-class | Accuracy | 0.834 ± 0.0084 | 0.838 +/- 0.00677 |
| | Balanced Accuracy | 0.735 ± 0.0258 | 0.719 +/- 0.0248 |
| | MCC | 0.793 ± 0.0105 | 0.797 +/- 0.00831 |
| | ROC AUC* | 0.969 ± 0.0026 | 0.977 +/- 0.00227 |
| | Ave. Precision Score* | 0.900 ± 0.0073 | 0.918 +/- 0.00392 |

**Figure 2:**
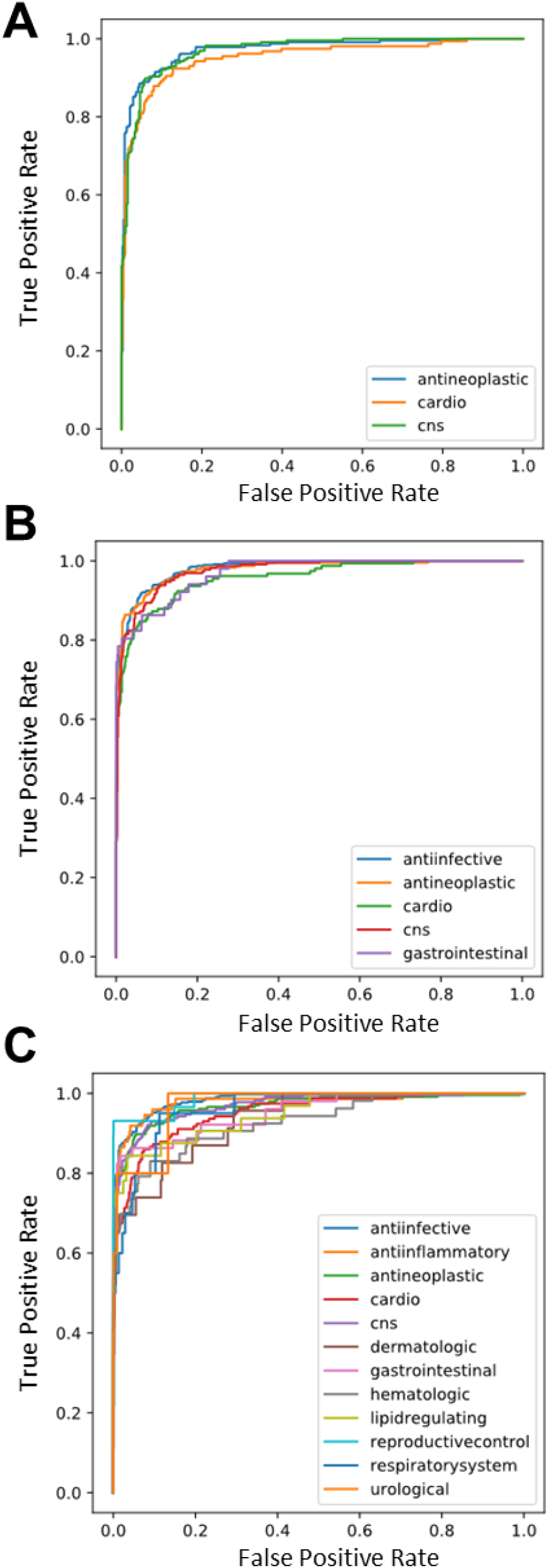
Receiver Operator Characteristic Curves from the IMG+CNN model predictions on the fifth validation set of the (A) 3-, (B) 5- and (C) 12-class datasets. Performance on this example fold is representative of the performance on all five folds shown in Table 4.

### Model and Representation Comparisons

Given the unequal stratification among examples within classes in the training and validation sets, the per-class performance of both models was compared on one representative validation fold. Confusion matrices of true class versus predicted class from IMG+CNN for the 3 subtasks reveal differences in per-class validation accuracy. In the 3-class prediction subtask, the prediction performance has similar high accuracy among the three groups. The most difficult class to predict is cardiovascular drugs; 13% of cardiovascular drugs are predicted incorrectly as central nervous system drugs (**Figure 3A**). In the 5-class prediction subtask, which includes the 3-class drugs plus gastrointestinal and anti-infective drugs, prediction performance is generally lower relative to the 3-class performance. The prediction accuracy typically follows the number of examples available with anti-infective drugs predicted at high accuracy (93%). Gastrointestinal drugs are the smallest class but are predicted more accurately (78%) than cardiovascular drugs (76%), which are still often confused for CNS agents (**Figure 3B**). In the 12-class prediction task, the accuracy for anti-infective molecules remains the highest (92%), and smaller classes are generally predicted less accurately (**Figure 3C**). The smallest class ‘Urological Agent’, which contains only 26 molecules, was rarely predicted correctly in the validation set (40%). The difference in class sizes likely contributes to this deficiency, and we did not directly control for this during training.

**Figure 3:**
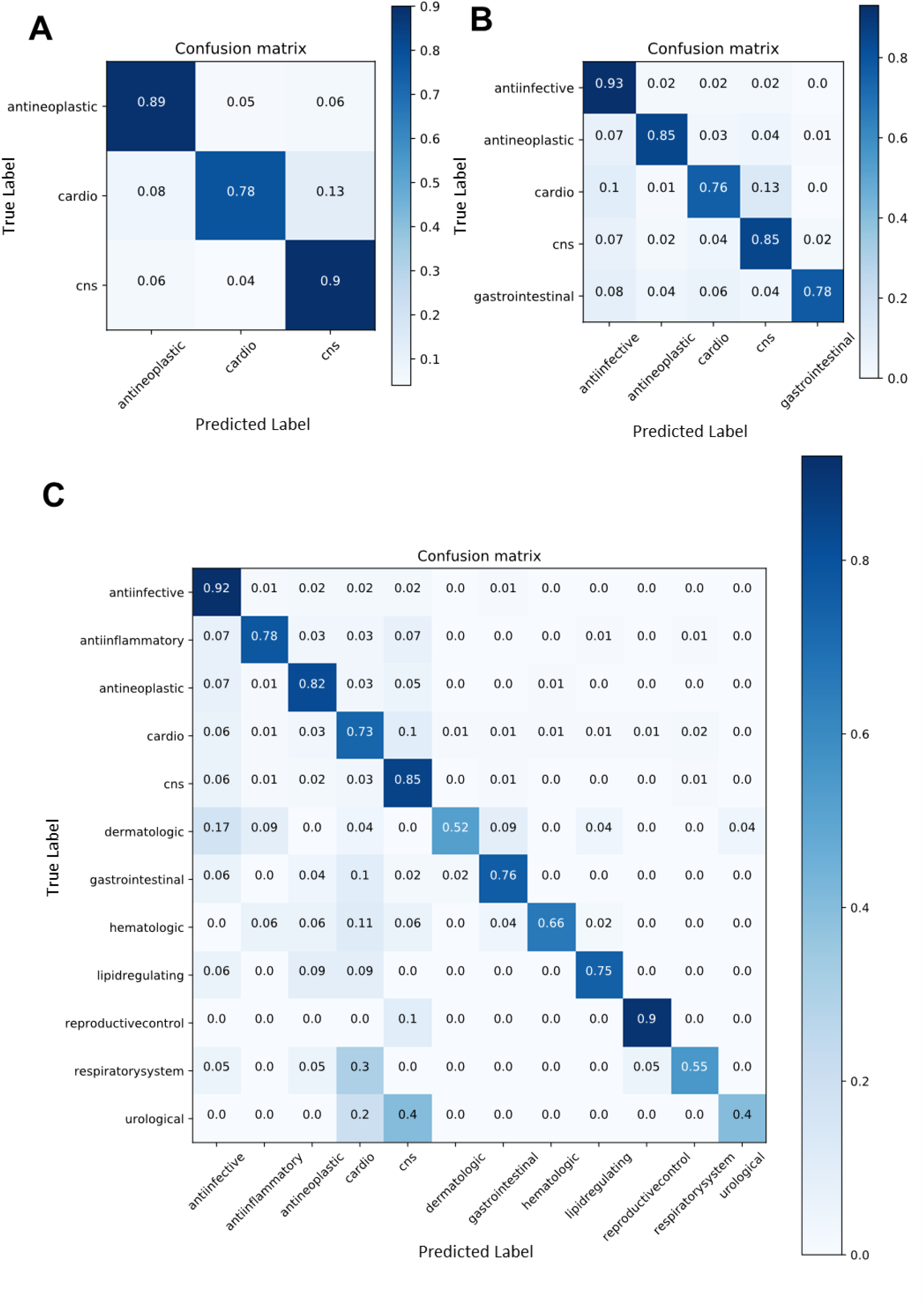
Confusion matrices from IMG+CNN classifiers from validation sets of drug molecules belonging to (A) 3-, (B) 5-, and (C) 12-classes sets of MeSH ‘Therapeutic Uses’. Each matrix shows the predictions from the fifth validation set using models trained on the large dataset.

The same analysis of per-class validation set accuracy for results from MFP+RF produced accuracy for each class within 5 percentage points but revealed a trend for different errors (**Figure S4**). MFP+RF more often over-predicted drugs as anti-infective, which may explain the 97% accuracy for that class. For example, the RF and CNN predicted 11% and 0%, respectively, of hematological drugs as anti-infective. There were also class-specific performance differences. MFP+RF models were better at predicting CNS agents correctly (MFP+RF: 90% vs IMG+CNN: 85%) and dermatologic agents (MFP+RF: 57% vs IMG+CNN: 52%), but the IMG+CNN models were better at predicting cardiovascular agents (MFP+RF: 68% vs IMG+CNN: 73%).

### Chemical Insight into Learned Molecule Properties

A compound’s drug-like properties are related to its chemical features such as molecular weight, lipophilicity, and the number of hydrogen bond donors and acceptors (e.g. quantitative structure activity relationships or QSAR). Lipinski’s rule of five famously states that a drug-like compounds should have no more than five hydrogen bonds donors, 10 hydrogen bond acceptors, a logP greater than 5 (related to hydrophobicity), and molecular weight under 500^3^. These properties are directly or indirectly encoded in the image- and fingerprint-based representations of chemical structure we use to train models.

To make inferences about what our models may have learned about chemical properties, we computed molecular weight, XlogP, and hydrogen bond donors and acceptors for our all single-class molecules and compared their distributions with one-way ANOVA and Games-Howell posthoc testing (**Figure 4, Figure S5, Table S2, Table S4**). Our IMG+CNN model often confused respiratory drugs with cardiovascular drugs (30%, **Figure 3**), and the chemical property analysis revealed that this drug class was indistinguishable from cardiovascular drugs with regard to the four computed properties (**Table S4** row 51, adjusted p-value = 1). The similarity of these properties may explain the confusion. Conversely, respiratory drugs are indistinguishable from gastrointestinal drugs across Lipinski’s properties, but both models can easily distinguish these two classes. This suggests that there are important structural features of drugs learned by the classifiers that fall beyond the conventional framework of how chemists understand drug-like chemical properties.

**Figure 4:**
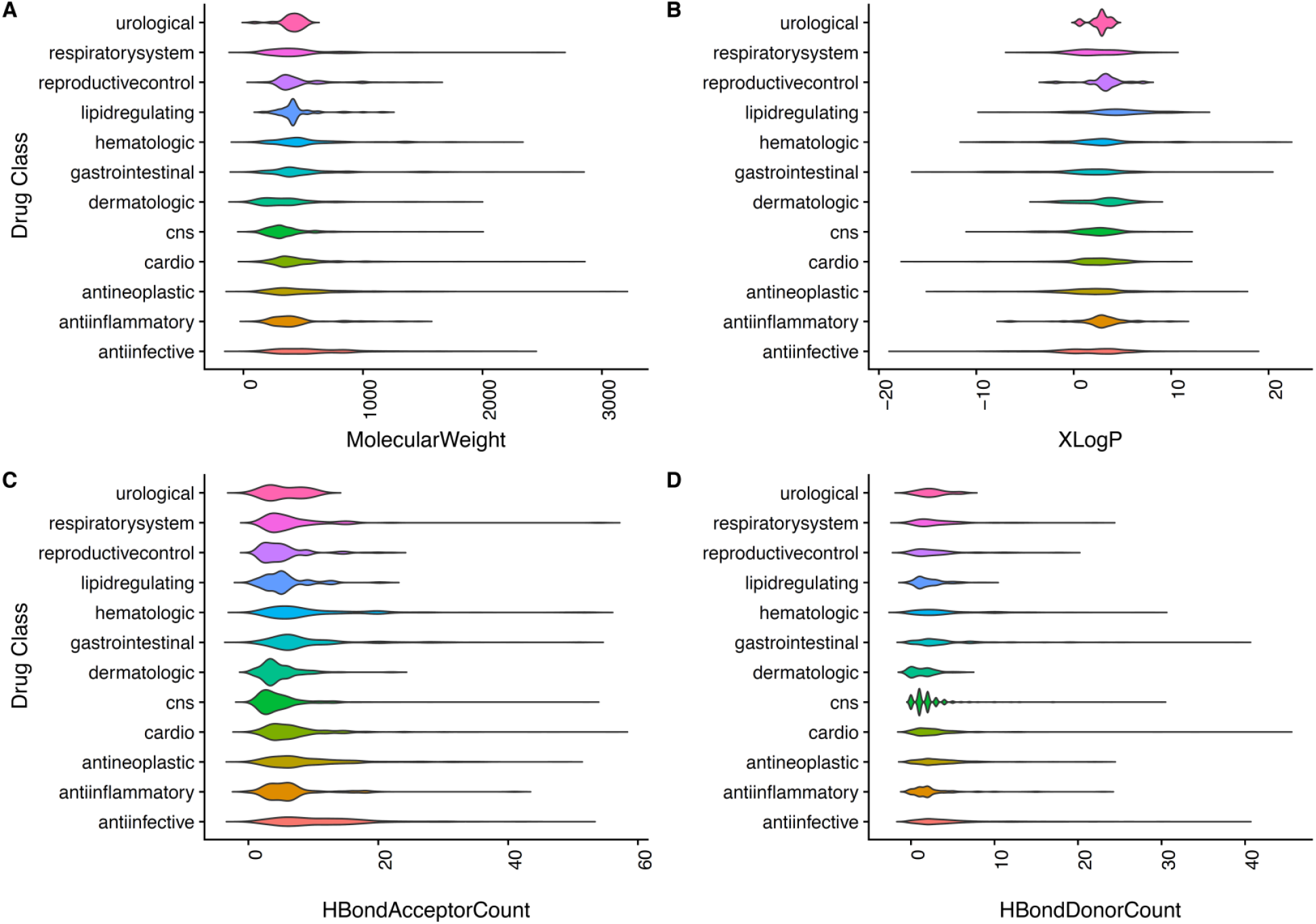
Class-level distribution of Lipinski’s drug-like properties: (A) molecular weight, (B) XlogP, (C) hydrogen bond acceptor count, and (D) hydrogen bond donor count.

However, other cases are less clear. Dermatologic drugs were often confused for anti-infective (17%) and gastrointestinal drugs (9%) despite all properties showing very significant differences (adjusted p-values < 3.6E-3). However, the MFP+RF model often confused dermatologic drugs for CNS agents (**Figure S4**), which is a mistake that the IMG+CNN model never makes (0%). This pair of chemical classes is statistically different in only the # of hydrogen bond donors. Thus, it is possible that the RF is underweighting this difference while the IMG+CNN model has learned to use it.

### Misclassification for Mechanism or Drug Repurposing Opportunities

Misclassification of drugs can be interpreted in at least two ways: (1) the model hasn’t learned enough to accurately predict the true class, or (2) the model has learned something new about the drugs and classes. Although the latter is more interesting, the former is the safer and more likely interpretation. However, cases where the model is wrong might present opportunities for drug repurposing. In addition, we hypothesize that those incorrect predictions might be useful for understanding drug mechanisms. For example, among the 6 molecules in the urological drug validation set, the IMG+CNN model misclassified Trospium as a central nervous system (CNS) agent. This is not surprising, however, because Trospium is known mechanistically as a muscarinic antagonist^57^, which is a common function of CNS drugs. In fact, the structure of Trospium is remarkably similar to another muscarinic antagonist used to treat Parkinson’s disease, Benztropine^58^ (**Figure 5**).

**Figure 5:**
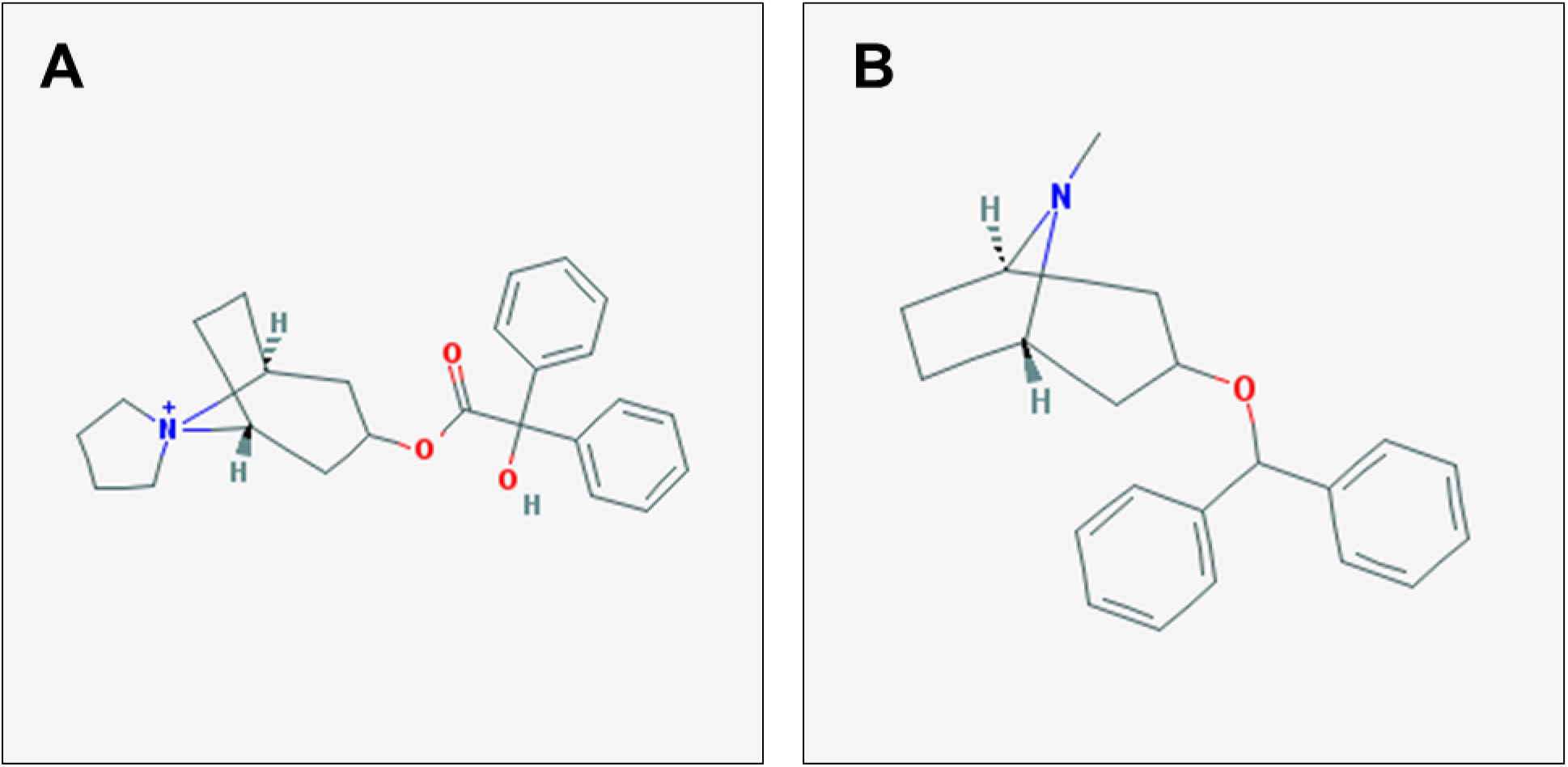
Example of misclassified drug that reveal mechanism and repurposing opportunities. (A) Structure of Trospium, a urological drug known to act as a cholinergic muscarinic antagonist, which was classified by the model as a CNS agent. (B) Structure of Benztropine, a muscarinic antagonist used to treat Parkinson’s disease.

### Multi-label Classification

Although we restricted analysis in above sections to only drugs with one annotated class, multi-label classification is an extension where we allow more than one drug class to be predicted for each example. When not filtered for only molecules in a single class, the set of all molecules with these 12 MeSH Therapeutic uses contains 8,336 molecules assigned a total of 9,885 classes. This corresponds to an average of 1.2 classes per molecule. We used a separate IMG+CNN model to learn these drug classes in the multi-label setting and evaluated the accuracy, F-beta score, ROC AUC, and average precision score (**Table 5**). The model achieved excellent performance in all computed metrics, but the ROC AUC and average precision scores show the multi-label prediction task is more challenging than the single-label version (**Table 4**).

**Table 5:** Multi-label classification of the 8,336 molecules matching 9,885 classes. Results are from 5-fold cross validation with folds determined by iterative stratification. Accuracy, ROC AUC, and average precision scores are not directly comparable to the 12-class single-class formulation (**Table 4**) because the number of molecules differs. *Class score thresholds set to 0.5.

| Problem group | Metric | IMG+CNN |
| --- | --- | --- |
| 12-class multi-label | Thresh. Accuracy* | 0.954 +/- 0.00133 |
|  | F-beta* | 0.635 +/- 0.0168 |
|  | ROC AUC | 0.938 +/- 0.00353 |
|  | Ave. Prec. Score | 0.837 +/- 0.00953 |

To better understand the multi-label prediction performance, true and predicted pairwise drug class memberships were used to generate drug class networks (**Figure 6**). The predicted and true network relationships were similar overall, but some connections were missing from the predicted network, such as between the anti-infective class and both the lipid regulating and hematological classes. Most of the true relationships were recovered in the predicted relationships, but the model did overestimate the strength of the most common relationships.

**Figure 6:**
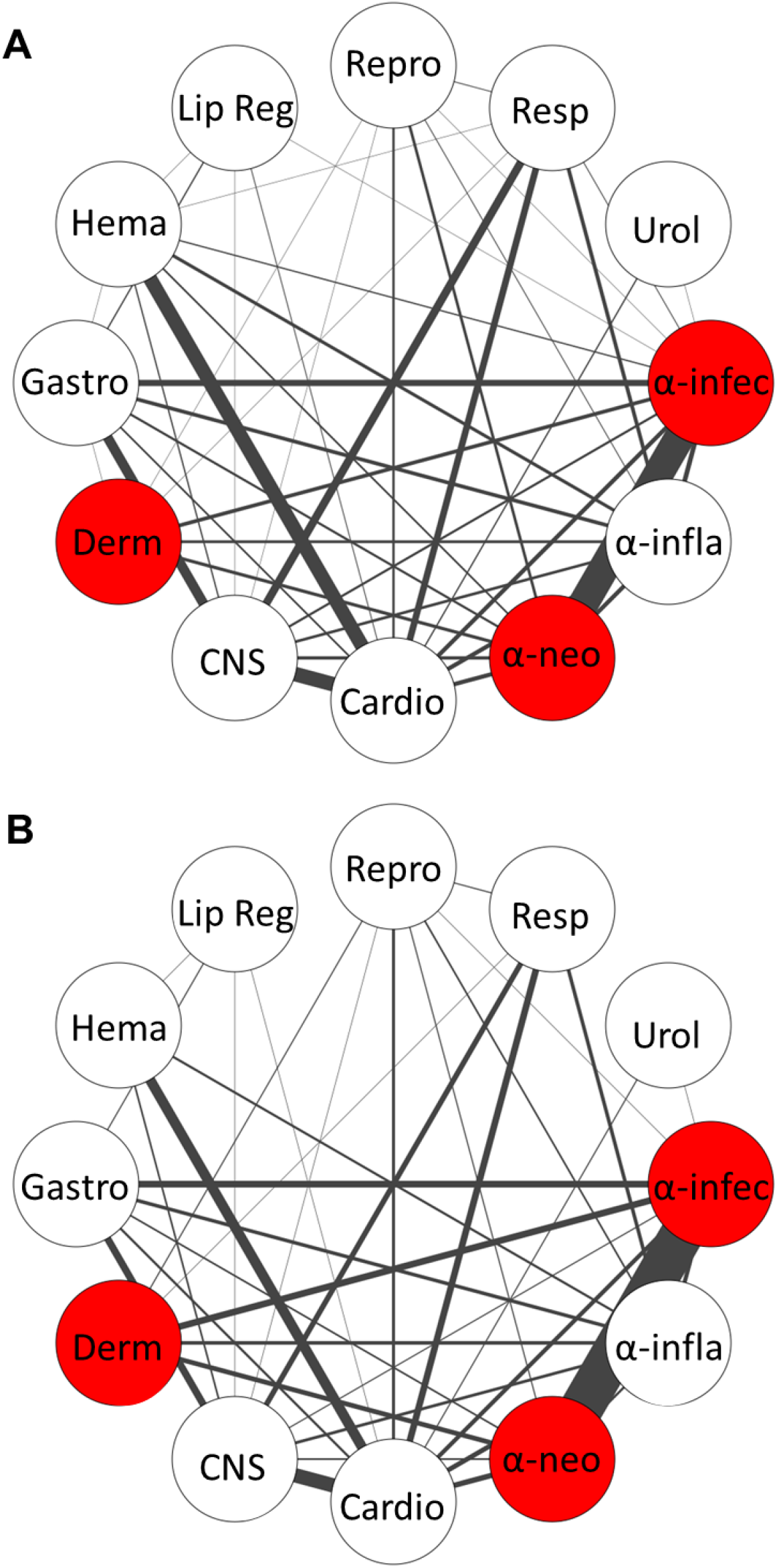
Analysis of multi-label drug classification using the IMG+CNN model. A network of relationships was computed from the (A) true class labels or (B) predicted class labels of the fifth validation fold. The width of each edge denotes the strength or frequency of co-occurrence. Red nodes indicate class grouping determined by their co-occurance.

## DISCUSSION

Here we report two drug classification models that greatly exceed a previous benchmark on the same prediction task. Our models use molecular structures directly as inputs, whereas the previous study used alterations of the transcriptome as a proxy for molecules. The results presented here suggest that experimental measurement of a molecule’s influence in biological systems may not be needed to accurately predict some types of chemical properties, such as annotated drug classes. However, we do not believe this necessarily means that direct empirical measure of the system is useless. Rather, additional research is required to determine which types of chemical prediction tasks require information about the biological state induced by chemicals and what type of biological state information is most useful (e.g. omic data, cellular morphology, etc.). There is likely room to improve effect-based models that would outperform molecule structure-derived models, especially on more complex prediction tasks.

Because the models presented here do not require empirical measurements of chemicals’ effects, they can be broadly applied to predict drug class after training; images and MFPs can be generated directly from the chemical structure for *any* chemical. A major limitation of the Aliper *et al.* featurization, or any empirically determined featurization, is that it requires new experiments for each new compound. This limited Aliper *et al.*’s total dataset to only 676 drugs for which transcriptomic data was available, thereby fundamentally limiting the utility of prediction to fewer compounds, and resulting in lower accuracy on that smaller dataset. Therefore, there must be substantial improvement in predictive performance to justify the extra cost of using experimentally-derived features in a virtual screening or chemical prediction setting. We propose that future studies on chemical prediction tasks that use empirically-determined featurizations also use models that consider only chemical structure features as a baseline.

Although there are several chemistry problems where DNNs outperform other shallow machine learning methods^49,59,60^, here the MFP+RF performed best with the small dataset of 676 molecules in the 5- and 12-class predictions. However, in the 3-class task with the small dataset, and all the tasks with the large dataset, the two models produced accuracies that were nearly indistinguishable. Because the performance of our two models was similar on the larger dataset, our results suggest that the CNN has more difficulty learning many classes from a small amount of data. This highlights that in general, more complex models should be benchmarked against strong standard machine learning methods, especially when training data is limited.

Much can be learned about chemical function from the cases where we find misclassification of chemical structures. We show cases where this can be rationalized by chemical properties of the molecules and cases where these properties that we often use to define the character of a chemical cannot explain the classification performance.

In the latter case, this may mean that our models have learned something about chemistry that may not be recognized by chemists. Still, the class-specific differences in molecular properties are interesting to compare. Further, when the models misclassify a structure, we can interpret this both as suggestive of a shared drug mechanism, and as an opportunity for drug repurposing. Drug repurposing is an especially important aspect of this work because application of an already-approved drug is much less costly that *de novo* approval of a new chemical.

An extension of the idea that misclassification can be used for repurposing or side effect prediction is multi-label classification. Our initial experiments followed the setting from Aliper *et al.* that excluded chemicals with multiple therapeutic uses, but we also extend the concept to a multi-label prediction model. The results show that our strategy is effective for the more complex multi-class prediction and that true relationships between drug classes are learned and recovered even with a relatively small dataset of less than 10,000 molecules. This model may be useful in predicting off-target effects of drugs, or discovery of repurposing opportunities. Taken together, our multi-label classification results prove the feasibility of this strategy with relatively simple modern deep learning packages.

## Acknowledgements

This work was supported by grants from the NIH NIGMS (P41 GM108538 and R35 GM118110 to JJC). JGM was supported by an NIH T15 fellowship (T15 LM007359). SL was supported by the University of Wisconsin-Madison Office of the Vice Chancellor for Research and Graduate Education with funding from the Wisconsin Alumni Research Foundation. This research was performed using the compute resources and assistance of the UW-Madison Center for High Throughput Computing in the Department of Computer Sciences.

## Author Contributions

Conceptualization, JGM and AG; Methodology, JGM, SL, and AG; Software, JGM, SL, IJM, and AG; Validation, JGM and SL; Formal Analysis, JGM, SL, and IJM; Investigation, JGM, SL, IJM; Resources, JJC and AG; Data Curation, JGM; Writing – Original Draft, JGM; Writing – Review and Editing, JGM, AG; Visualization, JGM, SL, IJM; Supervision, JGM, JJC, AG; Project Administration, JGM and AG; Funding Acquisition, JGM, JJC, AG.

## Supporting Information Available

Table S1 – Nested cross validation results from the hyperparameter search for the IMG+CNN using the 12-class 676 molecule dataset.

Table S2 – Filtered set of molecules and their computed drug-like properties.

Table S3 – Proportion of molecules used for drug property calculation in each class with XlogP values available.

Table S4 – Adjusted p-values from the Games-Howell posthoc test comparing drug properties for each class with one-way ANOVA

Supporting Information – Supplementary Figures S1-S5

## Table of Contents Graphic

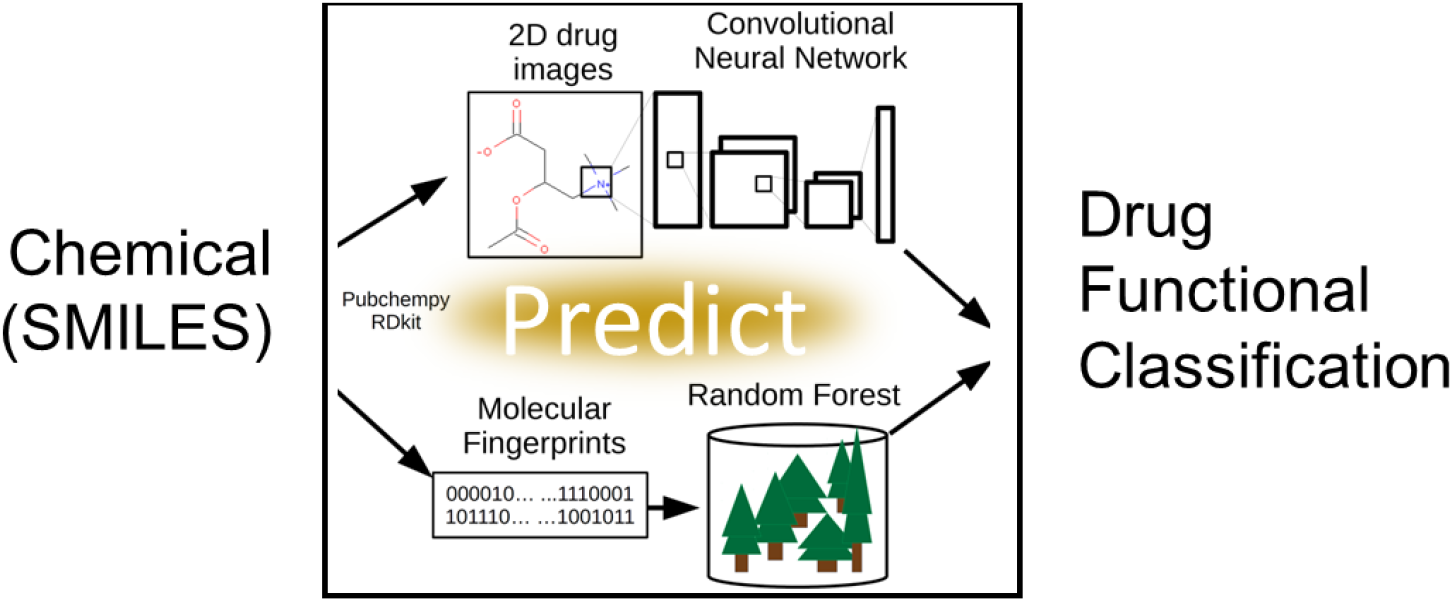

