## Supplementary Information for "Learning Drug Function from Chemical Structure with Convolutional Neural Networks and Random Forests"

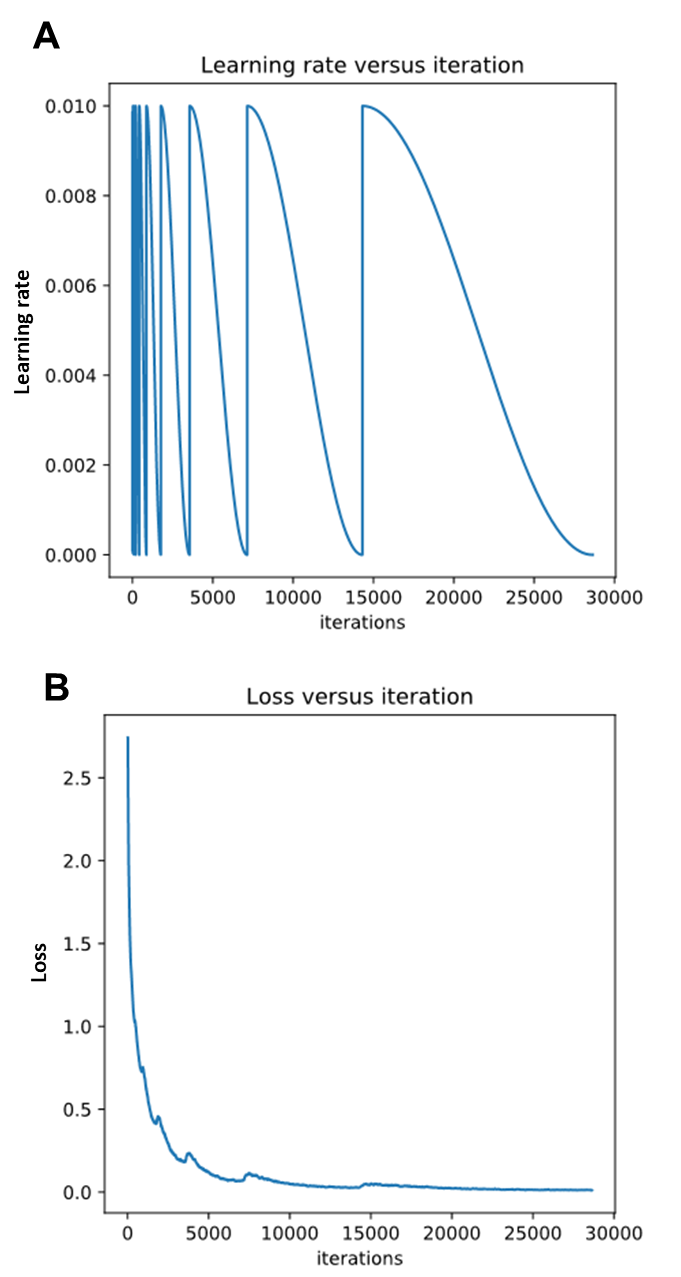

**Figure S1: Example of the cosine annealing learning rate strategy used for training the small and large single class models.** Seven cycles each with double the time for rate decay were used, resulting in a total of 127 epochs of training. (A) Learning rate versus training batch. (B) Loss versus training batch.

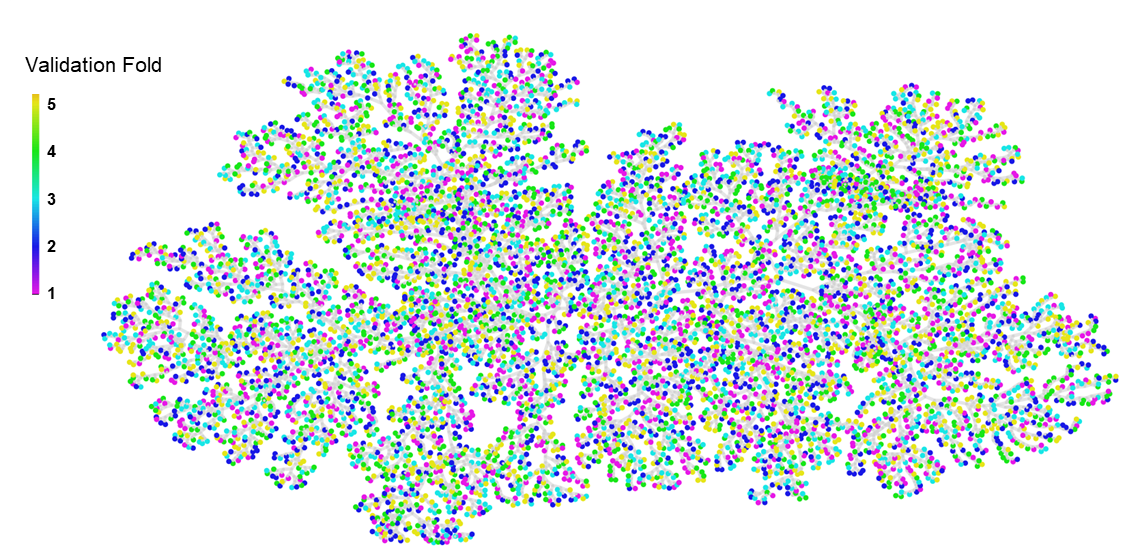

**Figure S2: ChemTreeMap showing random chemical similarity of the molecules in each of the 5 validation folds for the large dataset (6,955 molecules)**.

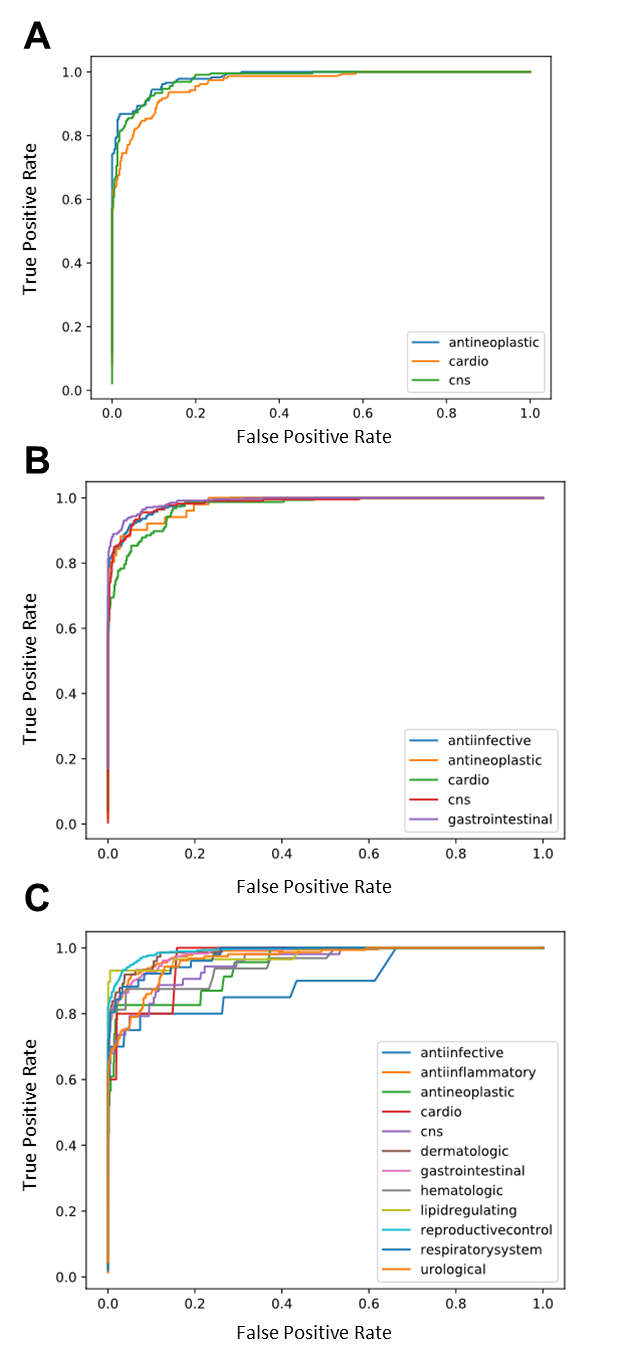

**Figure S3: Receiver Operator Characteristic Plots from the Random Forest Models trained with Morgan Molecular Fingerprints on the fifth validation set for the (A) 3, (B) 5, and (C) 12 therapeutic classes**.

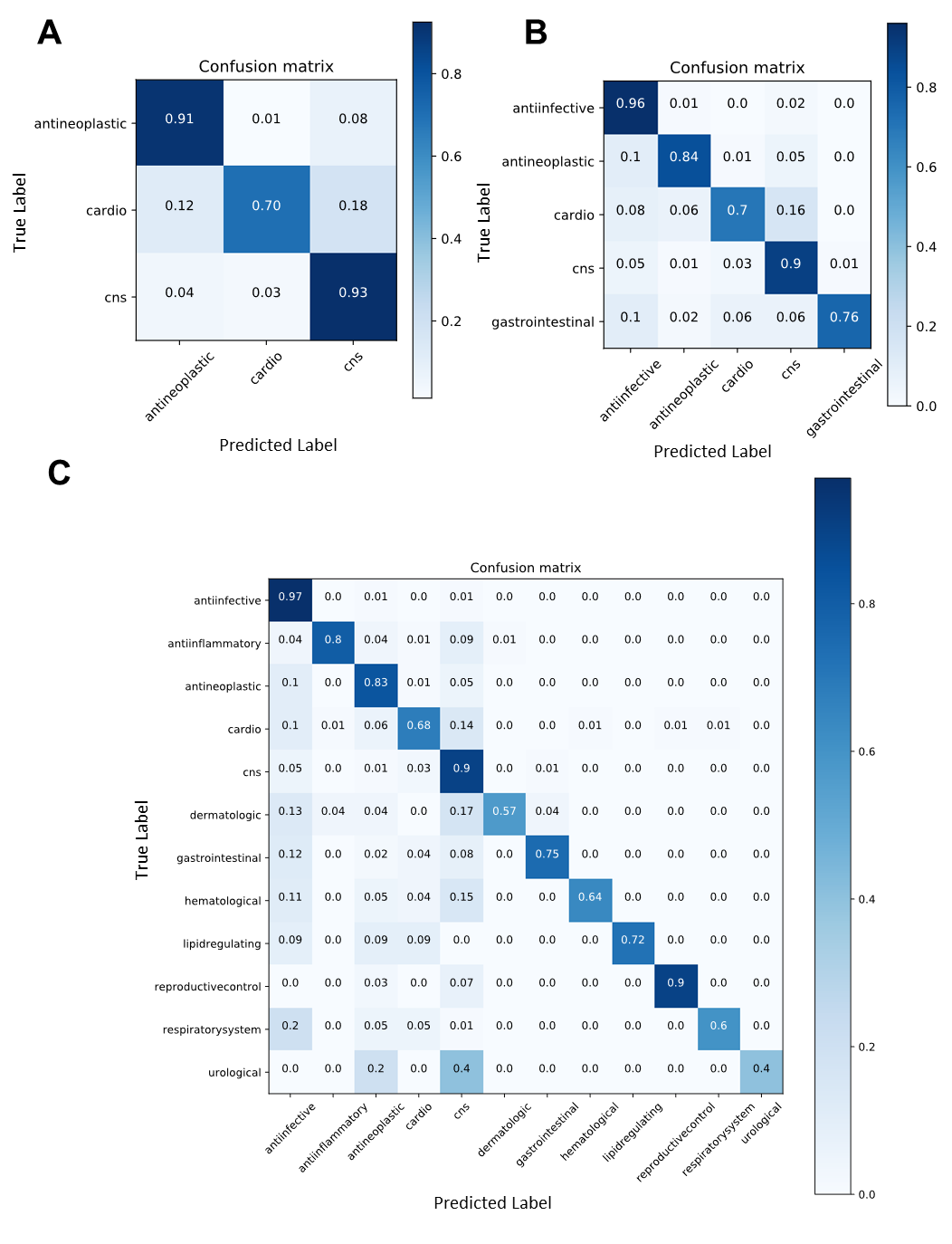

**Figure S4: Confusion matrices from MFP+RF classification performance over sets of drug molecules belonging to (A) 3, (B) 5, and (C) 12 therapeutic classes**. Each matrix shows the predictions from the fifth validation set using models trained on the large dataset.

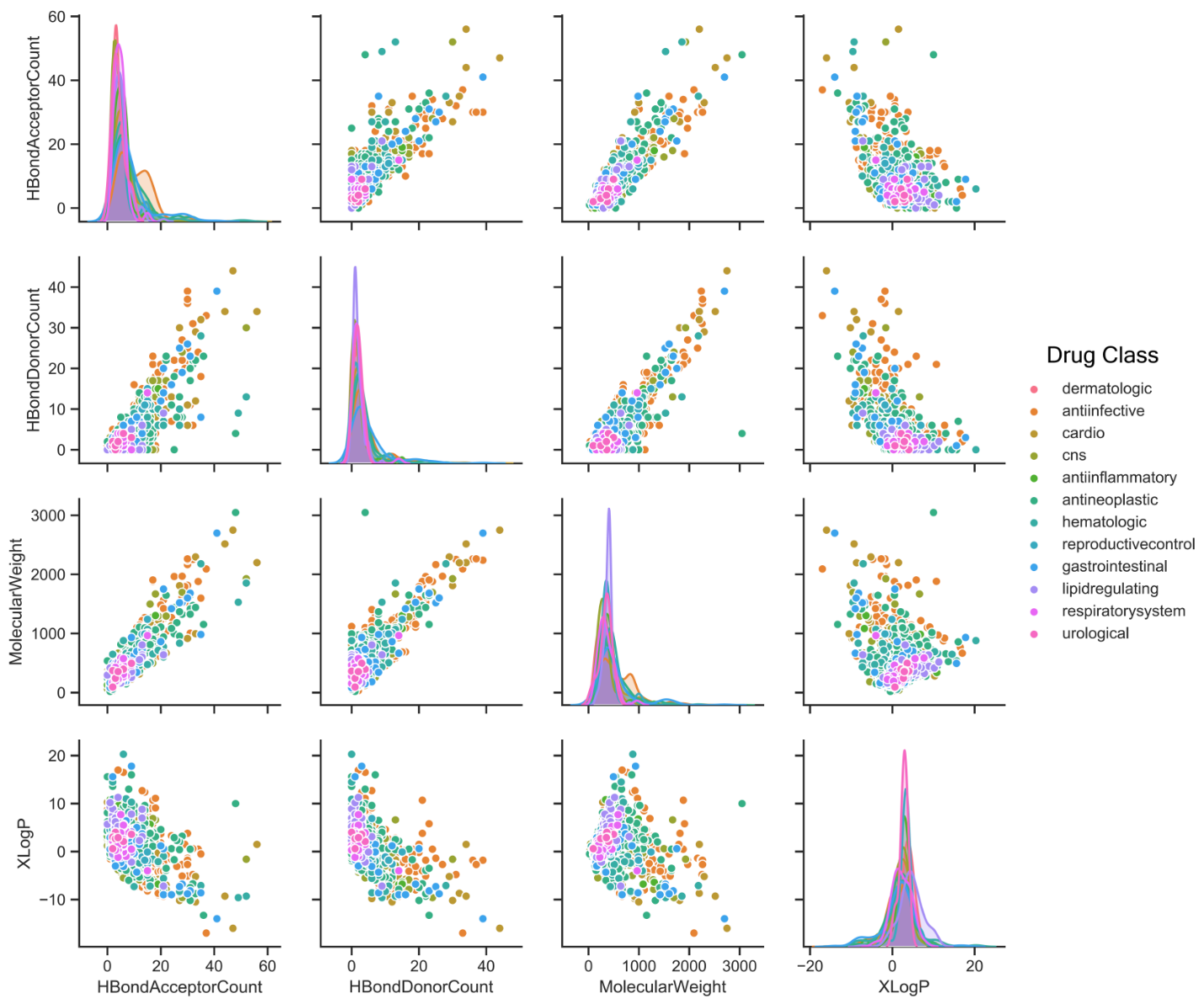

**Figure S5: Pairplots showing the distributions and correlations among the molecule properties for each drug classification.**
